# MetaOmixTools: a user-friendly web suite for meta-analysis of ranked features and functional enrichment

**DOI:** 10.64898/2026.02.24.707748

**Authors:** Rubén Grillo-Risco, Maksym Kupchyk Tiurin, Carla Perpiñá-Clérigues, Francisco J. Cordero Felipe, Samuel Lozano Juárez, Beatriz Dolader-Rabinad, Paula Palero-Renart, Silvia Salvador-Guerrero, Celine I García-Rodríguez, María de la Iglesia-Vayá, Francisco García-García

## Abstract

The growing number of omics datasets in public repositories provides an opportunity to enhance data reusability through data integration; however, complex statistical barriers often hinder the effective combination of independent studies. To address this problem, we present MetaOmixTools, an interactive web-based suite that streamlines the meta-analysis of ranked feature lists and functional enrichment profiles. The platform integrates two primary modules - MetaRank and MetaEnrich within a code-free environment. MetaRank generates robust consensus rankings from multiple lists by implementing weighted (e.g., Rank Product) and unweighted (e.g., Robust Rank Aggregation) strategies, while MetaEnrich performs functional meta-analyses by combining probability values from individual over-representation analyses using established statistical techniques. Using case studies, we established consensus rankings for acute spinal cord injury across heterogeneous platforms, identifying conserved inflammatory marker genes in the upregulated gene list (e.g., *Slpi, Ccl2, Msr1*) and synaptic loss genes in the downregulated gene list (e.g., *Kcna2, Dao, Ppp1r1b*), and also characterized inverse functional intersections between melanoma brain metastasis and neurodegenerative diseases. By providing intuitive, real-time visualization and reproducible workflows, MetaOmixTools empowers the research community to extract consistent biological insights from multi-study data. We have made MetaOmixTools freely available at https://bioinfo.cipf.es/metaomixtools/.

**Graphical Abstract:** 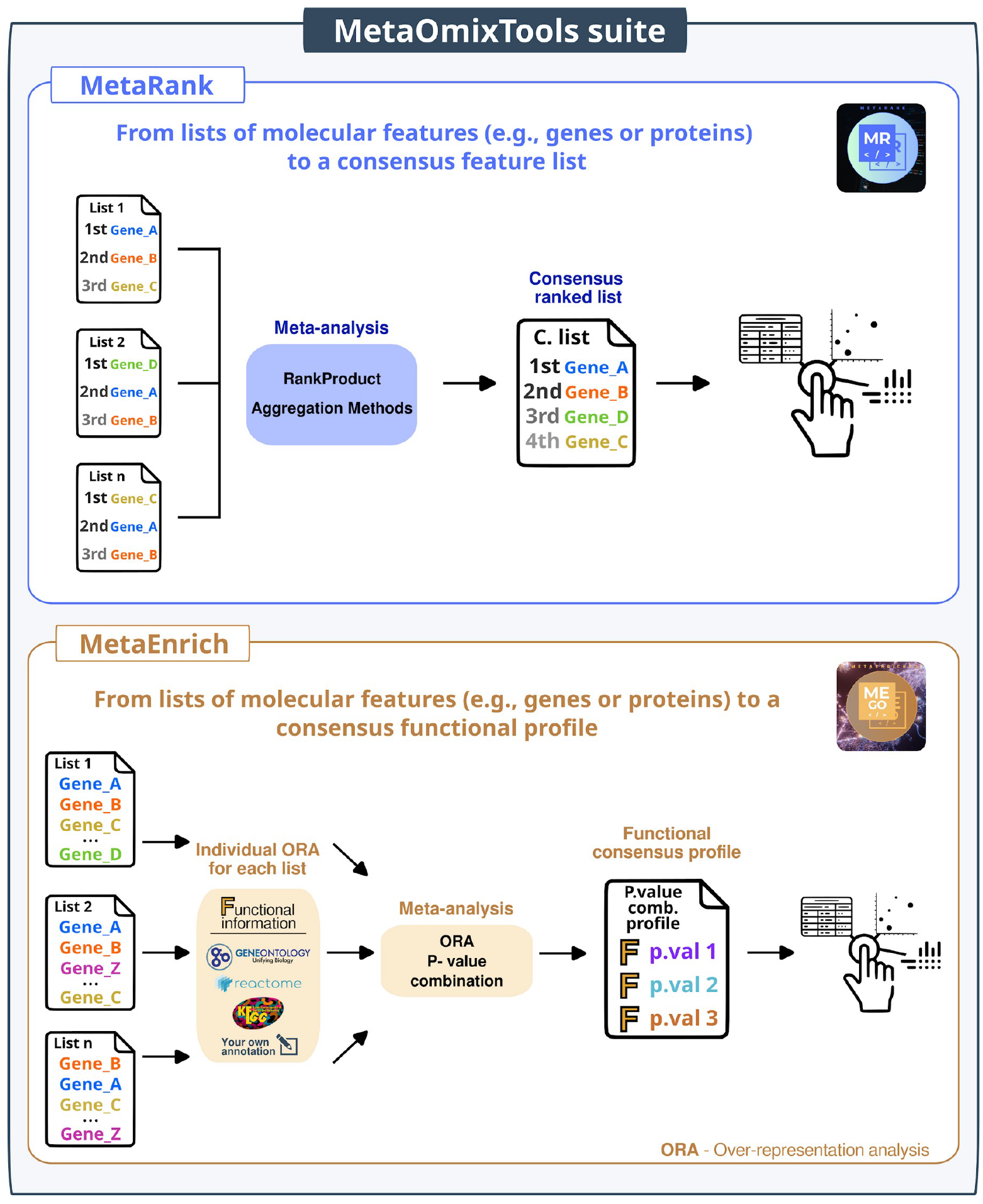

## BACKGROUND

High-throughput technologies have generated a rapidly increasing number of omics datasets for humans and model organisms. Public repositories such as Gene Expression Omnibus (GEO) and ArrayExpress contain thousands of studies that address similar biological questions [1,2]. Combining results across multiple datasets significantly increases reproducibility and statistical power, particularly in situations where individual studies remain small or have been conducted under heterogeneous experimental conditions [3–5]. Two meta-analysis strategies hold special relevance for this purpose: i) integrating ranked feature lists to produce a single consensus ranking and ii) combining p-values from functional enrichment analyses to create a consensus functional profile.

Rank aggregation methods enable researchers to identify those genes, proteins, or other molecular entities consistently ranking at the top across lists from multiple sources; for example, these methods integrate differential gene expression analyses or prioritize genes from independent genomic variation studies using pathogenicity indicators [6–8]. This approach employs diverse methodologies, with the choice of method depending on the nature of the input data. The availability of quantitative scores allows the application of weighted strategies (e.g., log-fold changes or p-values), whereas unweighted approaches apply when only the rank order represents a known quality.

While various R packages exist in the literature [refs], RankProd and the Robust Rank Aggregation (RRA) algorithm remain the most widely adopted tools for weighted and unweighted approaches, respectively. Their widespread adoption, with more than 2,000 citations to date, provides evidence of the reliability and robustness of these methodologies.

Meta-analysis for functional enrichment has been applied since the expansion of omics data in the early 2010s to identify shared biological functions across multiple feature lists [9–12]; meta-analysis in this context is typically based on p-value combination methods. The metap R package implements several standard statistical methods for this purpose (e.g., Fisher’s, Stouffer’s, Tippett’s, and Wilkinson’s methods) [13–16], and is actively maintained on The Comprehensive R Archive Network (https://cran.r-project.org). The thousands of monthly downloads reflect the continued utility of this package in statistical and bioinformatics workflows.

Although R packages contain these methodologies, a reliance on programming remains a significant barrier for many researchers. Web-based tools have been developed to make these approaches more accessible. However, existing platforms often present limitations such as a restricted range of meta-analysis methods, inflexible input formats, or obsolescence due to lack of maintenance [9,17,18]. Consequently, there is a need for intuitive, well-maintained platforms that facilitate access to meta-analysis methods without compromising statistical rigor and providing flexible statistical strategies.

To bridge this gap, we introduce MetaOmixTools as an open-access, registration-free web application that integrates the MetaRank and MetaEnrich frameworks. The platform offers an interactive interface that guides users through the entire workflow, from data upload to result interpretation, generating customizable, real-time visualizations. By eliminating the need for coding expertise, MetaOmixTools democratizes ranking-based and p-value-based meta-analysis, making sophisticated data integration accessible to bioinformaticians and experimental biologists alike.

## METHODS

The application structure is modular, comprising two primary analytical modules—MetaRank and MetaEnrich—implemented as independent components yet integrated into a unified user interface (**Figure 1**).

**Figure 1.**
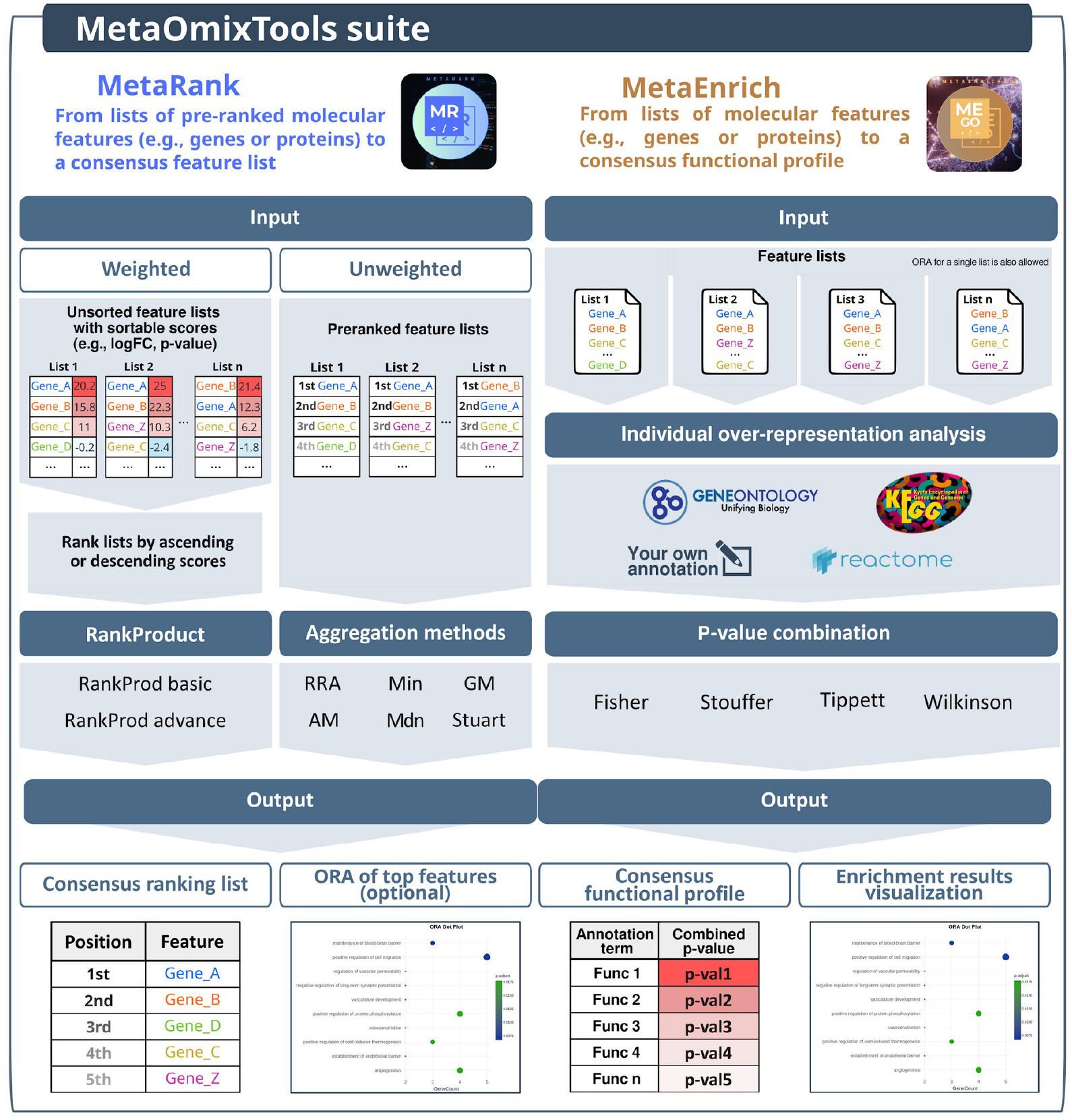
Overview of the MetaOmixTools suite workflow. The MetaRank module (left) integrates multiple lists of molecular features, provided either as unsorted lists with associated scores (weighted input) or as pre-ranked lists (unweighted input). In weighted mode, features are ordered by ascending or descending scores and combined using rank-based meta-analysis methods, including Rank Product (RP) (basic and advanced). The unweighted mode applies aggregation strategies such as Robust Rank Aggregation (RRA), minimum (Min), geometric mean (GM), arithmetic mean (AM), median (Mdn), and Stuart’s method. The main output is a consensus ranking list, with optional over-representation analysis (ORA) applied to top-ranked features. The MetaEnrich module (right) performs functional enrichment meta-analysis from multiple feature lists. ORA is first carried out individually for each list using annotation databases (Gene Ontology, Kyoto Encyclopedia of Genes and Genomes, Reactome) or user-defined annotations, and the resulting p-values are then combined across lists using methods (Fisher’s, Stouffer’s, Tippett’s, or Wilkinson’s) to generate a consensus functional profile. As a secondary option, individual ORAs can also be performed on a single feature list without the meta-ranking step. All outputs include interactive visualizations and downloadable tables and plots.

### MetaRank module

The MetaRank module, designed to obtain a robust consensus ranking from multiple feature lists (e.g., genes or proteins) to identify consistently relevant elements across studies, implements two distinct statistical approaches depending on the nature of the input data (**Figure 1**, left panel). Weighted lists containing quantitative metrics (e.g., fold changes or p-values) implement the Rank Product (RP) method via the RankProd package [19]. This non-parametric approach calculates the geometric mean of ranks for each feature across all replicates and assesses significance against a null distribution generated by permutation, yielding a percentage of false prediction (PFP) score equivalent to false discovery rate (FDR). For unweighted pre-ranked lists, the module integrates the RRA algorithm using the RobustRankAggreg package [20]. RRA uses a probabilistic model based on order statistics to detect features ranked consistently better than expected under a random null model. Alternative aggregation methods, such as geometric mean or median rank, are also available for specific research needs. Finally, to facilitate biological interpretation, the module allows users to perform an optional over-representation analysis (ORA) on the top-ranked features of the generated consensus list.

### MetaEnrich module

The MetaEnrich module performs functional meta-analysis by combining enrichment results across multiple feature lists. The workflow consists of two sequential statistical steps (**Figure 1**, right panel). First, an ORA is executed independently for each input feature list using the hypergeometric test implemented in the *enricher* function of the clusterProfiler package [21]. Features are mapped to biological terms using internal local databases (Gene Ontology, Kyoto Encyclopedia of Genes and Genomes [KEGG], or Reactome) for *Homo sapiens, Mus musculus*, and *Rattus norvegicus*, or via custom user-provided annotations. Second, the module statistically combines the p-values obtained for identical functional terms across different lists to derive a global consensus significance. The metap package [22] was used to implement several classic p-value combination methods, enabling users to tailor the analysis to their specific biological hypotheses. These options include Fisher’s (sum of logs), Stouffer’s (sum of z-scores), Tippett’s (minimum p-value), and Wilkinson’s (maximization of p-values) methods [13–16]. The final consensus p-values are adjusted for multiple hypothesis testing using the Benjamini-Hochberg method [23]. Beyond multi-list meta-analysis, the module offers the flexibility to analyze a single feature list, serving as a conventional ORA tool without integration.

### Implementation and user interface

MetaOmixTools was developed using R (v4.3.3) and deployed as an interactive web application using the Shiny framework (v1.10), utilizing HTML and CSS for enhanced front-end interactivity and responsiveness (**Figure 2**). The application architecture is modular, with each module encapsulating its own user interface logic, backend R scripts, test datasets, and graphical resources, while sharing core utility functions for data processing.

**Figure 2.**
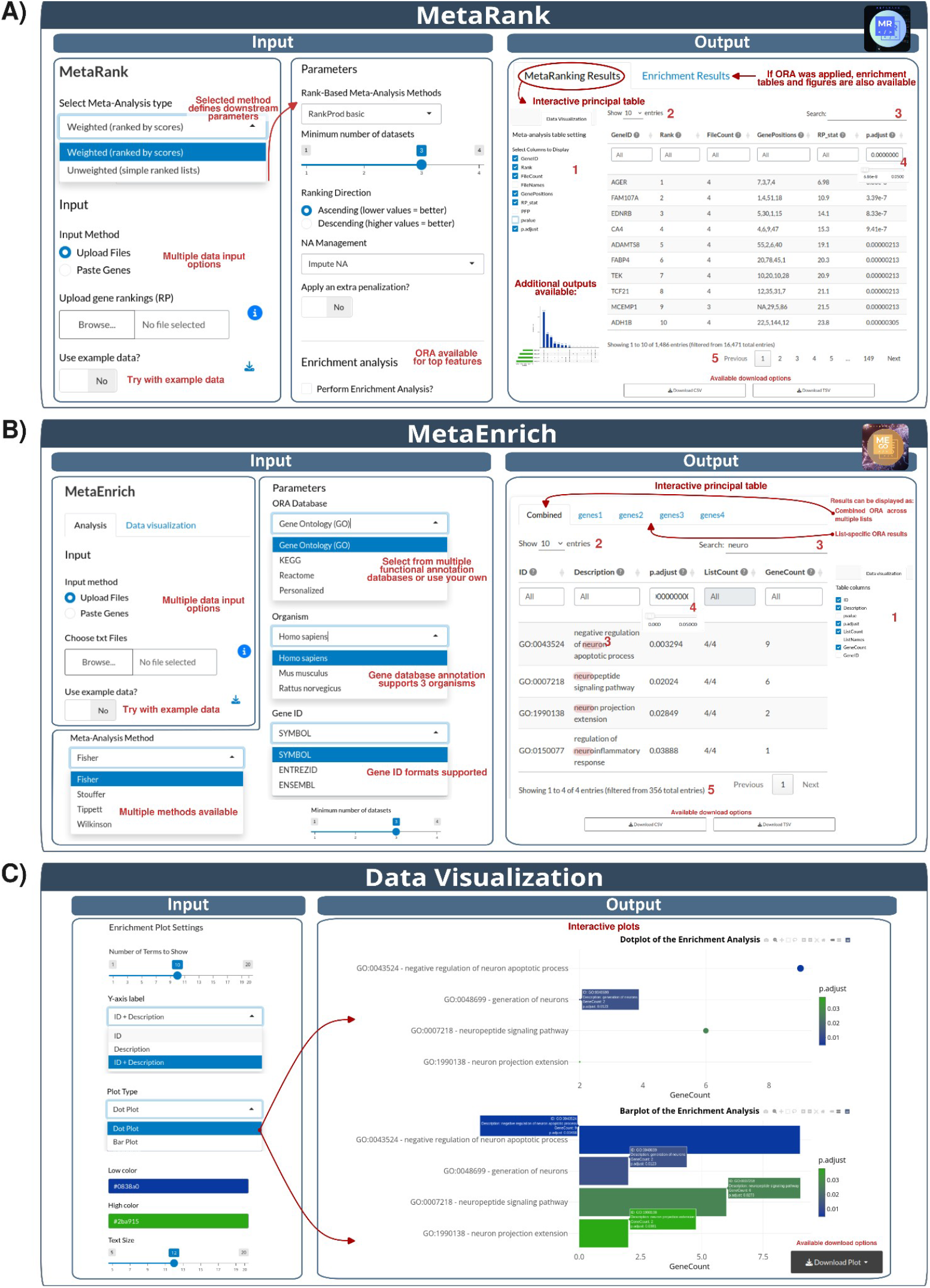
MetaOmixTool suite interface. Input and output options of the MetaRank and MetaEnrich modules and data visualization panel are shown. Multiple data input options are supported, including file upload, manual gene input, and execution with example data. All result tables and plots can be downloaded for further analysis or reporting. **A**. MetaRank module. The selected meta-analysis strategy automatically defines downstream parameters. Over-representation analysis (ORA) can be optionally applied to top-ranked features. Results are displayed as an interactive meta-ranking table, with additional secondary outputs available. **B**. MetaEnrich module. Functional enrichment analysis is performed across multiple feature lists using different annotation databases, supporting three organisms and multiple gene identifier formats. Uploading custom annotations is also possible. Results can be visualized as combined enrichment profiles or as list-specific ORA results. **C**. Data visualization. ORA results for both modules allow interactive visualization using dot or bar plots, with customizable display parameters (e.g., number of terms, axis labels, color gradients, and text size). Numbers indicate standard DT table options for searching, filtering, pagination, and column-specific queries.

To facilitate ease of use, both modules share standardized input and output mechanisms. Data can be introduced via multiple file uploads, direct pasting of feature lists, or by loading built-in example datasets. Similarly, outputs are harmonized across the platform. Tabular results are displayed using interactive DT tables [24] with filtering, sorting, and export options (**Figure 2A and Figure 2B**).

The MetaRank interface allows users to easily toggle between weighted and unweighted meta-analysis modes; crucially, the interface provides specific mechanisms for handling missing identifiers across heterogeneous datasets. Users can choose to impute missing ranks with the median rank of the list, ignore incomplete cases, or penalize missing features by assigning them the worst possible rank, ensuring that the final consensus reflects global consistency across studies (**Figure 2A**).

The MetaEnrich interface allows users to select the p-value combination method. The incorporation of local, lightweight annotation databases across major model organisms is a critical feature of this design, ensuring full functionality and speed without reliance on external application programming interfaces (API) during execution. Furthermore, users can also upload custom files. Additionally, the interface includes filtering options, such as defining the minimum appearance frequency of a feature across lists to be considered in the combination, allowing for stricter consensus definitions (**Figure 2B**).

The visualization engine represents a core component of the user experience (**Figure 2C**). The Enrichment Analysis Plot panel renders results from both modules using dynamic graphics implemented with ggplot2 [25] and plotly [26]. Users can visualize consensus results using dot or bar plots, which are fully customizable in real time. Parameters such as color gradients, axis labels, number of terms shown, and text sizes can be adjusted instantly, allowing users to export publication-quality figures directly from the browser.

## RESULTS

To illustrate the utility of MetaOmixTools, we applied two case studies to demonstrate the critical functions of the MetaRank and MetaEnrich modules.

### Case study 1. MetaRank module: Obtaining consensus metarankings from differential expression analysis results

To demonstrate the capability of MetaRank to integrate heterogeneous gene lists, we downloaded publicly available differential expression results reported in the Meta-SCI app (https://metasci-cbl.shinyapps.io/metaSCI/) [27] derived from four independent GEO datasets (GSE464, GSE45006, GSE104317, and GSE133093) that compared moderate acute spinal cord injury (SCI) against control tissue. The selected datasets represent outputs from distinct experimental platforms, including the Affymetrix microarray (Rat Genome U34 Array and Rat Genome 230 2.0 Array) and the Illumina NextSeq 500 RNA-seq platform; this diversity provides a representative sample of heterogeneous transcriptomic sources suitable for meta-analysis. Overall, we aimed to establish a consensus ranking of genes consistently deregulated following acute SCI.

For each dataset, we separated genes into two lists according to their log-fold change (logFC) - upregulated (logFC > 0) and downregulated (logFC < 0) – and then uploaded these lists into MetaRank as independent inputs. We selected the weighted metaranking option, ranking genes based on their logFC values (**Supplementary File 1**), and computed two separate consensus rankings (for upregulated and downregulated genes) using the RankProd Basic method. A minimum of three occurrences across datasets was required for a gene to be included in the meta-analysis, given that GSE464 contains fewer probes than the other platforms. We set the ranking direction to descending (higher values = better) for upregulated genes and ascending (lower values = better) for downregulated genes; meanwhile, we ignored missing genes without penalization. Finally, we performed functional enrichment analysis of the top 200-ranked genes using Gene Ontology Biological Process (GO BP) annotations.

**Figure 3A** displays the top 10-ranked genes for the upregulated and downregulated consensus lists. Among the top upregulated genes, we found well-known inflammatory mediators, such as *Slpi* (Secretory Leukocyte Peptidase Inhibitor), *Ccl2* (C-C Motif Chemokine Ligand 2), and *Msr1* (Macrophage Scavenger Receptor 1), supporting their established role in immune activation after SCI [28–30]. Conversely, the top downregulated genes included critical markers of neuronal homeostasis and excitability, such as *Kcna2* (Potassium Voltage-Gated Channel Subfamily A Member 2), *Ppp1r1b* (Protein Phosphatase 1 Regulatory Inhibitor Subunit 1B), and *Dao* (D-Amino Acid Oxidase), reflecting the extensive repression of neuronal signaling and loss of tissue integrity typical of the acute injury phase [31–33].

**Figure 3.**
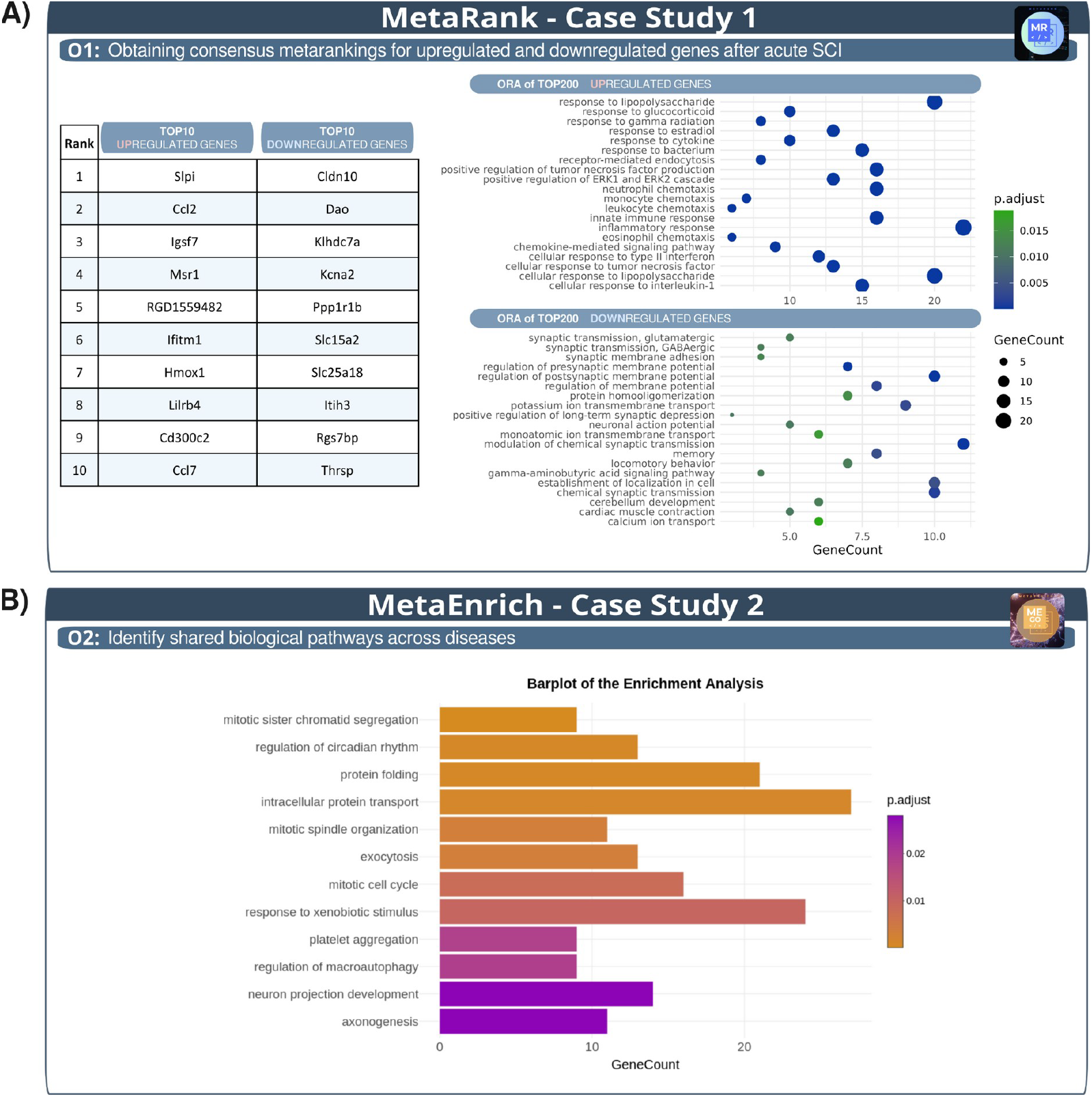
Validation of MetaOmixTools Through Biological Case Studies. **A**. MetaRank - Case Study 1. The table depicts the top 10-ranked genes for the upregulated and downregulated consensus lists. The associated dot plots illustrate distinct functional profiles following overrepresentation analysis of the top 200 genes in each consensus ranking. **B**. MetaEnrich - Case Study 2. The bar plot displays significantly enriched biological processes identified by p-value combination meta-analysis.

To interpret the biological significance of the consensus lists, we performed ORA on the top 200 genes from each ranking. Dot plots highlighted distinct functional profiles: we observed an enrichment of in immune-related processes (**Figure 3A**), such as *inflammatory response, chemokine-mediated signaling pathway, monocyte chemotaxis*, or *leukocyte chemotaxis*, in the upregulated metaranking but the significant depletion of neuronal functions, such as *potassium ion transmembrane transport, synaptic transmission, glutamatergic and locomotory behavior*, in the downregulated consensus signature, which agrees with the findings reported by Grillo-Risco et al. (2025) [27].

This case study illustrates how MetaRank can effectively integrate multiple transcriptomic data to produce robust consensus gene rankings and highlight biologically meaningful patterns across multi-study datasets.

### Case study 2. MetaEnrich module: Identifying shared functional programs across divergent transcriptional patterns in melanoma and neurodegenerative disease

To illustrate the ability of MetaEnrich to integrate functional evidence across multiple heterogeneous gene lists, we analyzed publicly available results from the MetaFun-MBM resource (https://bioinfo.cipf.es/metafun-mbm/) [34]. Shared molecular mechanisms exist between cancer and neurodegenerative diseases, as evidenced by the secretion of Alzheimer’s-associated amyloid β by melanoma cells and the repeatedly observed correlation between melanoma and Parkinson’s disease [35–39]. To explore these connections through functional meta-analysis, we selected four differential expression comparisons between disease samples and their respective healthy controls: melanoma brain metastasis (MBM-3), Alzheimer’s disease in hippocampus (AD-HP), Alzheimer’s disease in cortex (AD-CT), and Parkinson’s disease in substantia nigra (PD-SN). The neurodegenerative disease signatures derive from meta-analyses [40,41], whereas MBM-3 corresponds to a single independent study [42]. Therefore, these datasets remain suitable for the MetaEnrich evaluation under realistic multi-study heterogeneity. We aimed to capture the common biological pathways affected in these diseases by analyzing functions associated with divergent gene expression patterns: genes upregulated in cancer but downregulated in neurodegeneration.

To obtain the gene lists of interest, we first filtered for significant genes (FDR < 0.05) in all four cases. We selected the top 300 most significantly upregulated genes for melanoma and the top 300 most significantly downregulated genes for each of the three neurodegenerative datasets **(Supplementary file 2)**. We uploaded the four gene lists as input to MetaEnrich, then performed individual ORAs for each list using GO BP as the annotation database, and applied Fisher’s method to combine p-values, which required the evaluation of a functional term in all four lists before being considered for the p-value combination step.

The analysis revealed 12 significantly enriched BPs (FDR < 0.05) (**Figure 3B**), including processes related to cell cycle regulation (e.g., *mitotic sister chromatid segregation, mitotic spindle organisation, mitotic cell cycle*). The upregulation of these processes in melanoma correlates with the cancer-associated hallmark of sustained proliferation [43]; conversely, their downregulation in neurodegenerative diseases reflects the post-mitotic nature of neurons, where aberrant cell cycle re-entry often represents a prelude to neuronal apoptosis rather than division, or indicates the loss of proliferative glial support [38,44,45]. Finally, we observed terms related to neuronal structural development (e.g., *axonogenesis, neuron projection development*). The downregulation of these pathways in Alzheimer’s disease and Parkinson’s disease represents a direct molecular signature of neurodegeneration and synaptic loss [46,47], while their upregulation in melanoma highlights the “neural mimicry” of these tumors. Melanoma cells, which share an embryological origin with neurons, reactivate such neural developmental programs to acquire high plasticity and invasive capabilities, facilitating metastasis to the brain [34,35].

This case study illustrates how MetaEnrich can effectively integrate gene lists of diverse origins to identify robust consensus functional terms. As demonstrated in this case by the identification of shared functional processes in melanoma and neurodegenerative diseases, the tool excels at highlighting biologically meaningful patterns from heterogeneous data sources.

## DISCUSSION

Here, we present MetaOmixTools, a web-based suite that facilitates the widespread adoption of meta-analysis methods and makes them accessible to the broader research community. The platform integrates two modules—MetaRank and MetaEnrich—that allow users to generate consensus rankings and perform functional meta-analysis through p-value combination, respectively. Analyses can be carried out entirely online in an intuitive, interactive, and code-free environment, quickly producing results that can be easily explored and interpreted by experts and non-experts alike.

Versatility regarding input formats and statistical strategies represents a critical advantage of MetaOmixTools over existing solutions. While other tools require users to upload strictly preformatted or pre-ordered lists, we designed our suite to adapt to user data. We allow researchers to paste feature lists or upload raw files in standard formats (e.g., CSV or TSV). Notably, the platform automatically sorts these inputs based on user-defined scores (e.g., logFC in ascending or descending order), saving time and significantly reducing the risk of manual formatting errors common in data preprocessing. Furthermore, the suite offers precise control over data heterogeneity; users can easily define the minimum frequency of appearance for a feature to be included in the consensus, ensuring consistency across studies. In parallel, MetaRank provides specific mechanisms to handle missing values (e.g., imputation, exclusion, or penalization), guaranteeing that the final consensus reflects biological trends rather than technical artifacts from incomplete datasets. This flexibility also extends to the statistical methodology; while comparable tools often limit unweighted analysis to only three algorithms, MetaRank implements six distinct aggregation methods and offers basic and advanced modes for weighted RP analysis [17]. Similarly, MetaEnrich outperforms platforms that rely solely on Stouffer’s Z-score by providing a comprehensive range of p-value combination strategies, including the widely used Fisher’s method [18].

Although initially conceived for gene-based analyses, we designed MetaOmixTools as fully extensible to any biological feature. While the suite includes built-in functional annotations for model species (human, mouse and rat) and widely used databases (GO, KEGG, and Reactome), users also can upload custom annotation files to perform enrichment analyses with their own feature identifiers, making the suite adaptable to a diverse range of organisms and biological elements (e.g. proteins, metabolites, or microRNAs). This flexibility expands the potential applicability of MetaOmixTools across different research contexts. Another strong point of the suite is that MetaEnrich can also work with single lists, functioning as a conventional enrichment tool alongside its main multi-list meta-analysis functionality, providing users with an extra level of versatility within the same interface.

Finally, we designed MetaOmixTools with a strong emphasis on usability and publication-ready outputs. Unlike many other web tools that offer static visualizations, the suite generates interactive, highly customizable graphics (content and style), allowing users to modify colors, labels, and layouts to create publication-ready figures. Critically, we address the common lack of support in web-based tools by integrating an extensive help section, detailed tutorials, and contextual help messages throughout the whole analysis pipeline.

In summary, MetaOmixTools provides a unified, accessible solution for ranked-list and functional-enrichment meta-analysis. By combining flexibility, interactivity, and methodological robustness, MetaOmixTools empowers the bioinformatics community to integrate multi-omics datasets and extract meaningful biological insights in a simple and reproducible way.

## Key Points

- MetaOmixTools is an interactive, web-based platform that streamlines the meta-analysis of ranked feature lists and functional enrichment profiles, including two powerful analysis modules: MetaRank and MetaEnrich.
- MetaRank generates robust consensus rankings across studies using both weighted (e.g., Rank Product) and unweighted (e.g., Robust Rank Aggregation) methods.
- MetaEnrich performs functional meta-analysis by integrating p-values from independent over-representation analyses using established statistical approaches.
- The platform offers intuitive visualization and reproducible workflows, enabling consistent extraction of biological insights from multi-study omics data.

## DECLARATIONS

### Ethics approval and consent to participate

Not applicable.

### Consent for publication

Not applicable.

## Acknowledgments

The authors thank the Principe Felipe Research Center (CIPF) for providing access to the cluster, co-funded by European Regional Development Funds (FEDER) to the Valencian Community 2014-2020. The authors also thank Stuart P. Atkinson for reviewing the manuscript.

## Data availability

The code is available at https://github.com/EMEKATE00/MetaOmixThttps://github.com/EMEKATE00/MetaOmixToolsools. The web tool is openly available at https://bioinfo.cipf.es/metaomixtools/.

## Supplementary data

Supplementary File S1. Input for the case study 1. MetaRank module: Obtaining consensus metarankings from differential expression analysis results.

Supplementary File S2. Input for the case study 2. MetaEnrich module: Identifying shared functional programs across divergent transcriptional patterns in melanoma and neurodegenerative disease.

